# Simulating drying and human impacts on river networks to evaluate biological quality indices performance through the lens of metacommunity theory

**DOI:** 10.64898/2025.12.01.691531

**Authors:** Zeynep Ersoy, David Cunillera-Montcusí, Martí Piñero-Fernández, Miguel Cañedo-Argüelles, María Mar Sánchez-Montoya, Núria Bonada, Núria Cid

## Abstract

1. Incorporating metacommunity perspectives into bioassessment represents a major challenge in the management of drying river networks, where drying-induced fragmentation compromises the performance of biological indices to assess their ecological status. Indeed, current indices focus on local community responses to stressors and neglect the effect of regional processes, such as spatiotemporal connectivity and dispersal, on metacommunity assembly.
2. In this work, we explored the effect of drying on the performance of a widely used biological index using metacommunity simulations on a synthetic drying river network. We assessed how different gradients of drying-driven fragmentation and human impact extent determine local richness and the biological index scores by combining simulations with biomonitoring information.
3. We used a coalescent metacommunity model to simulate the exchange of individuals between local communities along synthetic drying river networks. These networks were subjected to different drying extent, drying intensity and human impact extent scenarios. Additionally, we considered two major characteristics for each simulated taxon: (i) tolerance to human impacts and (ii) dispersal strategy (flying, swimming, or drifting).
4. For each simulation, we obtained local richness and the biological index value. Then, we calculated biological index performance, which we defined as the capacity to distinguish between impacted and non-impacted sites. Finally, we tested our approach in six non-impacted European drying river networks, from which drying information was available.
5. Our results showed that low spatiotemporal connectivity consistently led to decreased local richness and low scores of the biological index (reflecting poor biological quality). As drying extent and intensity increased, drying-induced fragmentation significantly reduced the biological index performance. For example, with a 50% increase in drying extent, index performance fell over 70% and at high drying levels, this performance dropped more than 90%. This decay followed a convex pattern, with a marked drop in performance as soon as drying appeared in the catchment and leveling off at higher drying extents.
6. *Synthesis and applications*: We show how simulations can be used to incorporate network fragmentation and metacommunity dynamics into the design and validation of biomonitoring tools, thereby increasing their efficiency.

## Introduction

River networks are increasingly subjected to drying due to global change pressures (Datry et al., 2023; Messager et al., 2021). Drying reshapes river networks at both spatial and temporal scales, affecting the number of reaches that contain water and how frequently they lose it. Together, these alterations reconfigure fluvial connectivity at the spatiotemporal level (Jacquet et al., 2022; Journiac et al., 2025). These changes concomitantly impact the exchange of individuals, species, and the flux of matter across river segments (Arias-Real et al., 2023; Cid et al., 2022; Datry et al., 2023; Sarremejane et al., 2021), altering community assembly and ecosystem functioning (Holyoak et al., 2020; Journiac et al., 2025; Marco Palamara et al., 2023). Although the importance of these regional-scale drivers has been acknowledged, their influence on river management remains far from clear, challenging the conservation of drying river networks against current global threats (Cid et al., 2020; Soria et al., 2020; Datry et al., 2023; Sarremejane et al., 2024).

River management practices, including biomonitoring, mostly focus on local-scale processes (i.e., environmental filtering), overlooking the key role that regional processes such as drying-driven fragmentation play in shaping diversity (Brodie et al., 2025; Chase et al., 2020; Cid et al., 2020; Heino et al., 2015). Consequently, traditional biomonitoring tools often fail to properly assess human impacts on drying river networks (Bonada et al., 2024; Ersoy et al., 2024; Soria et al., 2020). For instance, reaches that become isolated due to drying often have lower species richness despite favourable environmental conditions (Cañedo Argüelles et al., 2015; Henriques Silva et al., 2019). At the same time, highly connected sites may harbour greater richness even with suboptimal environmental conditions (Cañedo Argüelles et al., 2015; Heino, 2013; Jabot et al., 2020). In this context, metacommunity perspectives (Leibold & Chase, 2017) can provide key insights for river conservation by incorporating not only local drivers but also habitat position within the network (Borthagaray et al., 2020; Heino, 2013; Heino et al., 2015; Thompson et al., 2017) and the different dispersal strategies of organisms (Milošević et al., 2022; Siqueira et al., 2014; Stojković Piperac et al., 2023). Several works have highlighted the importance of these drivers in determining species richness and biomonitoring indices in perennial rivers (Durães et al., 2016; Siqueira et al., 2014). Nevertheless, this interplay remains unexplored in drying river networks due to the complexity of quantifying the spatiotemporal scales at which drying shapes connectivity and, in turn, alters metacommunity dynamics (Courtwright & Hawkins, 2024).

The link between river biomonitoring and metacommunity theory has been primarily explored from a theoretical point of view in perennial networks (Heino, 2013; Patrick et al., 2021). The few existing conceptual frameworks focused on drying river networks expect a decline in the performance of biological indices with increasing drying, which is modulated by spatial connectivity and organisms’ dispersal and resistance traits (Cid et al., 2020, 2022). Although considerable progress has been made in understanding how drying influences biodiversity from local to regional scales (Jacquet et al., 2022; Journiac et al., 2025), a mechanistic understanding of how drying may impact biological indices is still lacking. Within this context, the growing availability of hydrological data from drying river networks can be very useful, because it allows quantifying spatiotemporal connectivity (Cunillera Montcusí et al., 2023; Mimeau et al., 2024; Pineda-Morante et al., 2022). Indeed, several works have already highlighted the key role that spatiotemporal connectivity plays in drying river networks by determining community diversity and ecosystem functioning (Chalmandrier et al., 2025; Fernández Calero et al., 2024; Hárságyi et al., 2025). Thus, these advances open the window to assess the interplay between spatiotemporal connectivity and biomonitoring indices performance (Bonada et al., 2024; Cid et al., 2020).

In this study, we assessed the effects of river network drying and human impacts on the performance of a widely used biological index (Figure 1). To do this, we coupled macroinvertebrate biomonitoring databases with metacommunity simulations incorporating organisms’ dispersal across varying degrees of river drying (i.e., intensity and extent) and human impacts (i.e., extent). First, we investigated how local richness and the score of the biological index responded to the loss of spatiotemporal connectivity and widespread human impact extents. Second, we quantified the expected failure of biomonitoring along gradients of drying by calculating the performance of the biological index (i.e., the relative difference between impacted and non-impacted reaches). Finally, we assessed the potential changes in index performance for six European drying river networks from which long-term drying patterns were available. Building on previous works, we expected that *(i)* local richness and the score of the biological index would decrease with increasing drying and human impact extents (Escobar Camacho et al., 2025; Stubbington et al., 2024) and that *(ii)* the performance of the biological index would show a sharper decline when drying starts to fragment the river network (Courtwright & Hawkins, 2024). This drop would be driven by the loss in spatiotemporal connectivity and its interplay with human impacts, which would increasingly favour tolerant taxa. Finally, we expected that *(iii)* the decline if the performance of the biological index would attenuate as drying and human impacts affect most of the river network as a consequence of environmental (i.e., human impacts) and dispersal-driven metacommunity homogenisation (Piano et al., 2020; Rolls et al., 2023). Through this work, we provide useful insights for managers to determine at which level of drying the performance of biological indices might be compromised and fail to adequately detect human impacts in drying river networks.

**Figure 1.**
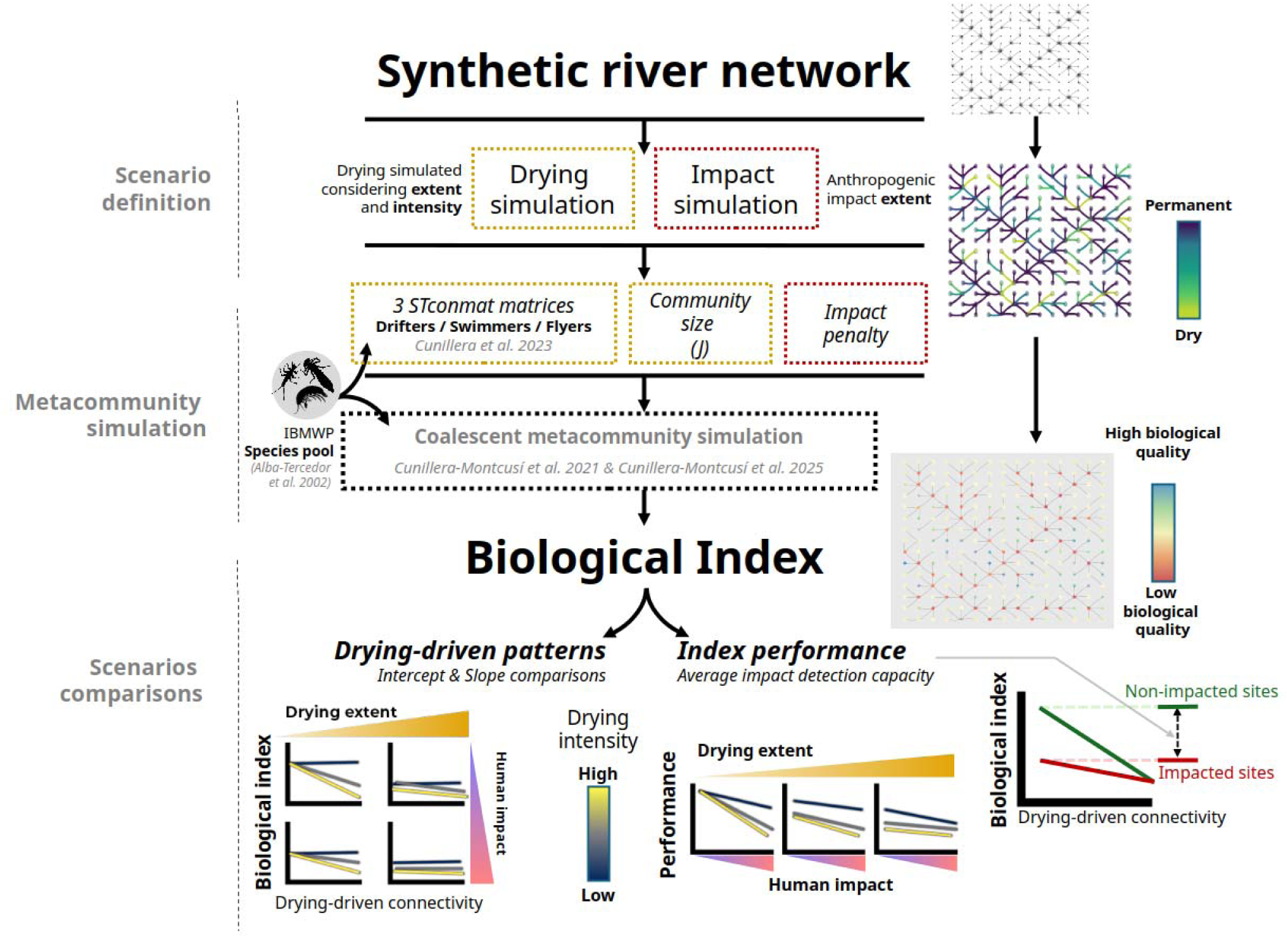
Overview showing the workflow to build the metacommunity simulations (Borthagaray et al., 2025; Cunillera Montcusí et al., 2021, 2023, 2025) and analyse their outputs to compare different scenarios.

## Material and Methods

### 1. Dendritic river networks

We generated a synthetic river network using the OCNet R package (Carraro et al., 2020), which represented our base template for simulating drying and human impact scenarios (Method sections 2 & 3). We used the *create_OCN* function to generate an Optimal Channel Network (OCN) to simulate a realistic river network on a square lattice (15 x 15) with a cell size of 2 km. This resulted in a network of 225 nodes representing a catchment of 900 km^2^ with a single outlet. This approach provides a standardized and replicable river network capturing the essential structure of a real mid-sized river, and ensures computational capacity for posterior simulations.

### 2. Drying scenarios and connectivity metrics

To consider the impacts of drying on the connectivity of the river network (i.e., probability of dispersal between nodes), we simulated a range of scenarios by manipulating two parameters: drying spatial extent (i.e., number of drying nodes throughout the network) and drying intensity (i.e., length and frequency of drying). Drying spatial extent indicated the number of dry nodes, ranging between 3 (1%) and 203 (90%) nodes of a total of 225. We defined nine levels of drying extent covering a range of drying intensity from almost permanent to fully intermittent networks (1, 10, 25, 35, 45, 50, 65, 75, 90%). Drying intensity represented the variability in duration and frequency of drying for each node, ranging from homogeneous and less frequent drying (e.g., all dry nodes remain dry for 10% of the time) to heterogeneous and highly recurrent drying (e.g., dry nodes are divided into different categories, ranging between nodes drying 10% and 99% of the time). We defined eight levels of drying intensity that summarise this gradient from low to high drying intensity (0.1, 0.25, 0.5, 0.55, 0.65, 0.75, 0.9, 0.99). Following this framework, we built a flow state database spanning 48 time units (i.e., a matrix of 225 columns corresponding to each network node and 48 rows with the flow state wet (1)/dry (0) at each time). This matrix corresponds to a 2-year time period, which represents a biologically relevant temporal window for these systems (Chalmandrier et al., 2025) and does not compromise computational times (Cunillera Montcusí et al., 2023). When selecting drying nodes, we always considered the same set of nodes to maintain comparability between drying scenarios of different spatial extents.

We calculated spatiotemporal connectivity matrices (STconmat) for each drying scenario, combining drying extent and intensity following Cunillera-Montcusí et al. (2023). STconmat described pairwise connectivity between all network nodes. We used a weighted network with distances as weights, which defines spatiotemporal connectivity as a proxy for dispersal resistance (*DirWei* scenario in Cunillera-Montcusí et al., 2023). Thus, greater STconmat values imply higher dispersal resistance (i.e., sites located far and/or isolated due to drying). Furthermore, we calculated STconmat for three different network structures corresponding to three major dispersal strategies extracted from Sarremejane et al. (2020) : aquatic passive (drifters), aquatic active (swimmers or crawlers), and aerial active dispersers (aquatic larvae with flying adults: flyers). For aquatic passive dispersers, we considered connectivity only from upstream to downstream. For aquatic active dispersers, we considered both upstream and downstream. For aerial active dispersers, we considered connectivity independently of the river network, only determined by geographical distance. Overall, STconmat values represented the potential for dispersal between any pair of sites by incorporating both distance-driven dispersal and network fragmentation caused by the defined drying scenarios.

### 3. Human impact scenarios

We also incorporated human impacts on metacommunity dynamics, defined as local environmental filters. We simulated two levels of impact at the local level: (i) Impacted, where we applied a penalty to establishment probability using tolerance scores from a standardised biological index based on macroinvertebrates at the family level in the Iberian Peninsula (IBMWP index, Alba-Tercedor et al., 2002; Appendix Table S1) (ii) Non-impacted, where establishment probability was solely determined by connectivity. Human impact extent was defined as the number of impacted nodes throughout the network. Although this biological index was developed for the rivers in the Iberian Peninsula, it is calibrated to other European indices (Munné & Prat, 2009), which allows its use as a general example for simulations. We explored several levels of human impact extent from 3 (1%) to 203 (90%) impacted nodes out of a total of 225 nodes and covering a gradient of different categories of impact extent (1, 10, 25, 35, 45, 50, 65, 75, 90%).

### 4. Metacommunity simulations in response to drying and human impact scenarios

We assessed how the interaction between drying and human impact influenced local biodiversity patterns by running a coalescent metacommunity model on the synthetic river network (Borthagaray, Cunillera-Montcusí, Bou, Biggs, et al., 2023; Borthagaray, Cunillera-Montcusí, Bou, Tornero, et al., 2023; Borthagaray et al., 2025; Cunillera Montcusí et al., 2021, 2025). This model simulated the exchange of individuals among nodes and along the river network, without accounting for biotic interactions (see Appendix S1). This exchange was determined by (i) probability of dispersal between nodes (i.e., STconmat) and (ii) human impact penalty for the impacted sites. Using this model, we simulated metacommunity assembly considering the constraints imposed by each scenario (i.e., drying extent, intensity, and human impact extent). For each metacommunity, we calculated total richness and biological index values for each node by adding family scores. We ran 10 replicates for each combination of scenarios and calculated their median to obtain an averaged value per scenario (see Appendix S1).

### 5. Data analysis

We explored the linear relationships between STconmat and two metrics: local richness and biological index across all scenario combinations (i.e., drying extent, and drying intensity, human impact extent). We extracted the intercepts and slopes of these relationships to quantify the trends of these two metrics along the simulated gradients. We then assessed the performance of the biological index, by using a reference scenario corresponding to a non-impacted river (Human impact extent = 0.01) with low drying extent (Dry extent = 0.01) and low drying intensity (Dry intensity = 0.1). Then, we calculated the performance of the biological index as the ratio of change between impacted sites and non-impacted sites relative to this reference as follows:

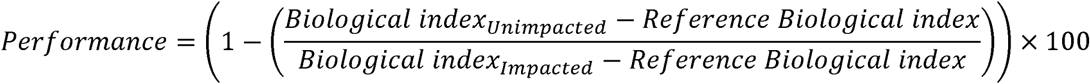

Performance values near 100% will imply that there is a maximal difference between impacted and non-impacted sites, indicating that the biological index can clearly detect impacts. The lower the % value, the lower the capacity of the index to distinguish between impacted and non-impacted sites.

For each of the defined drying scenarios (i.e., combination of drying extent and drying intensity), we fitted a general linear model between performance and anthropogenic impact extent to analyse how performance was impacted by drying along the anthropogenic impact gradient. Finally, we used the drying patterns from six non-impacted drying river networks located throughout Europe (Spain, France, Croatia, Finland, Hungary, and Czechia), from which daily drying information (site-level flow/dry state from January 2019 to 2021) was available at the catchment scale (Datry et al., 2021; Mimeau et al., 2024), to connect our simulations with real study cases. Details on each river network can be found at www.dryver.eu. Using this information, we quantified how the biological index performance would be impacted solely by drying in the river network, as these catchments are mostly non-impacted (anthropogenic impact set at 10%). We conducted all simulations and statistical analyses using R version 4.4.3 (R Core Team, 2025) and plotted graphs using the “ggplot2” and “viridis” packages (Wickham, 2009; Garnier et al., 2024).

## Results

### Simulation and metacommunity outputs

In total, we generated 648 unique combinations of drying extent, drying intensity, and human impact extent and simulated 10 different metacommunities for each combination (n = 6480 metacommunities). Overall, as drying extent and intensity increased, catchment-scale heterogeneity increased (Figure 2) with the concomitant decrease in perennial sites (Figure S1A) and more nodes presenting different drying frequencies (Figure S1B). These promoted regional-scale heterogeneity and shaped regional spatiotemporal connectivity along the different scenarios (Figure S1C).

**Figure 2.**
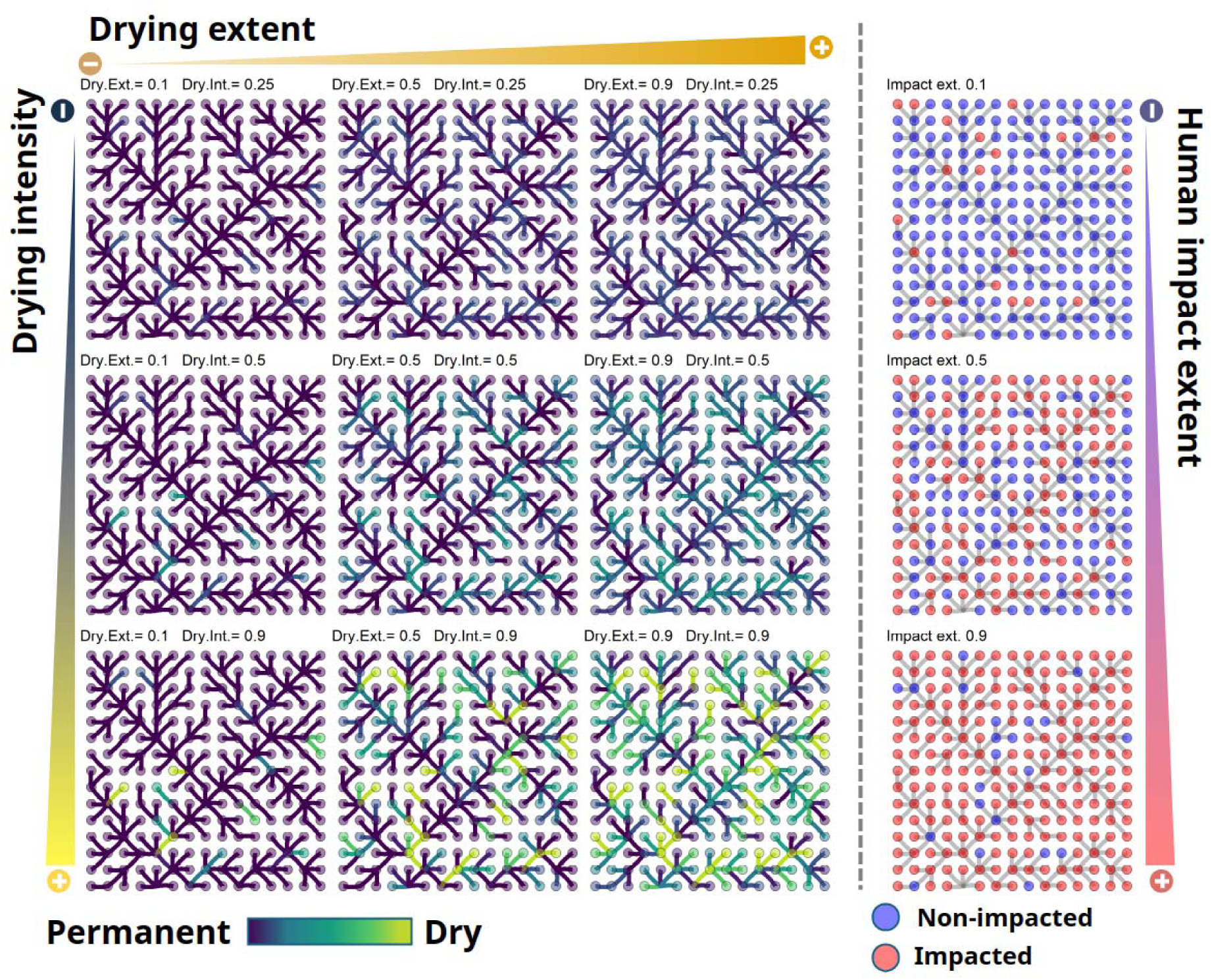
Examples of varying drying extent (Dry.Ext) and intensity (Dry.Int) and human impact extent scenarios (Impact.ext). The blue-green-yellow gradient illustrates the spatial configuration of water permanence across sites within the river network. The red and blue colours (left columns) indicate impacted and non-impacted sites, respectively.

### Response along drying-induced fragmentation

Both local richness and the score of the biological index decreased as spatiotemporal connectivity decreased (Figure 3, S2, S3). Thus, as a site became more isolated due to drying, its local richness and biological index scores decreased (Figure S4A). This pattern was clear under a low human impact extent (Figure 3, top row). As drying intensity increased, the slope of the relationship between the biological index and spatiotemporal connectivity increased, implying a stronger decrease in the scores of the biological index (i.e., the more frequently the river dried, the stronger its negative effect on index values; Figure 3 top row panels, S4B). However, as drying and/or human impact extent increased, these negative relationships disappeared and the effect of spatiotemporal connectivity faded (i.e., intercept decreased and slopes flattened; Figure 3 from top to bottom panels, S4B). This occurred because widespread drying and human impact extent resulted in low scores of the biological index across all sites, masking the negative effect of spatiotemporal connectivity (e.g., when all sites are greatly isolated and impacted, all sites present similar low index values). Therefore, as drying and human impact extents increased, all sites became equally taxa-poor, regardless of their connectivity (Figure 3, bottom panels).

**Figure 3.**
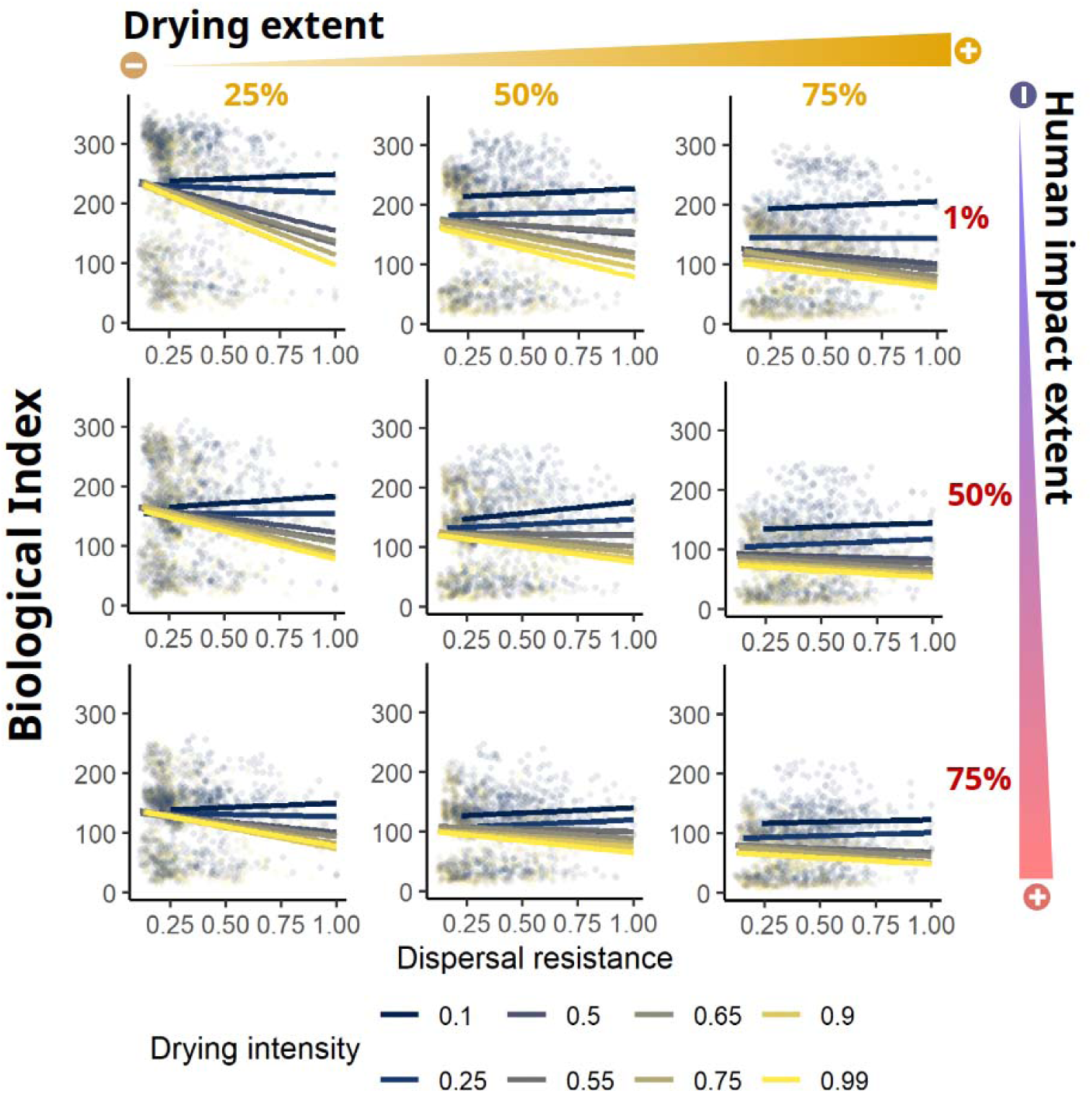
Linear relationships between the biological index and the dispersal resistance. Sites with high dispersal resistance are strongly isolated due to distance and drying (STconmat). Each panel corresponds to a combination of scenarios of drying extent (25 %, 40 %, 75 %), and human impact extent (1 %, 50 %, 75 %). For each combination, the considered drying intensities are represented as a linear trend (blue to yellow gradient).

### Biological index performance across scenarios

Drying-induced fragmentation and the greater extent of human impact resulted in a decrease in the performance of the biological index, as indicated by smaller intercept values (Figure 4, Figure S5). The negative slope between performance and human impact extent was modulated by drying extent and intensity (Figure 4, Figure S6). As drying extent increased, the decrease in intercepts suggested a reduced performance of the biological index (Figure 4). For example, under non-impacted conditions (i.e., 1 % human impact extent), an increase in drying extent from 10% to 25% reduced average performance by 26% (from 70% to 44%, respectively after averaging all drying intensities), while the same increase from 75% to 90% reduced it by 2.5% (from 9.5% to 7%, respectively after averaging all drying intensities). We observed a similar pattern with the increasing drying intensity. In scenarios with low drying extent, performance was only reduced by human impact extent (i.e. all lines decrease together, Figure 4, left panel). However, as both the drying extent and intensity increased, they overrode the effect of human impact by lowering the scores of the biological index across the network. As such, under non-impacted conditions (i.e., 1% of human impact extent) and 25% of drying extent, the difference between low and high drying intensity was around 27%, indicating the strong differentiation that drying intensity can also produce in intermediate scenarios. Under more extreme scenarios with high drying extent and intensity, the difference between impacted and non-impacted sites was consistently low, regardless of the degree of human impact extent (i.e., nearly flat slopes in Figure 4).

**Figure 4.**
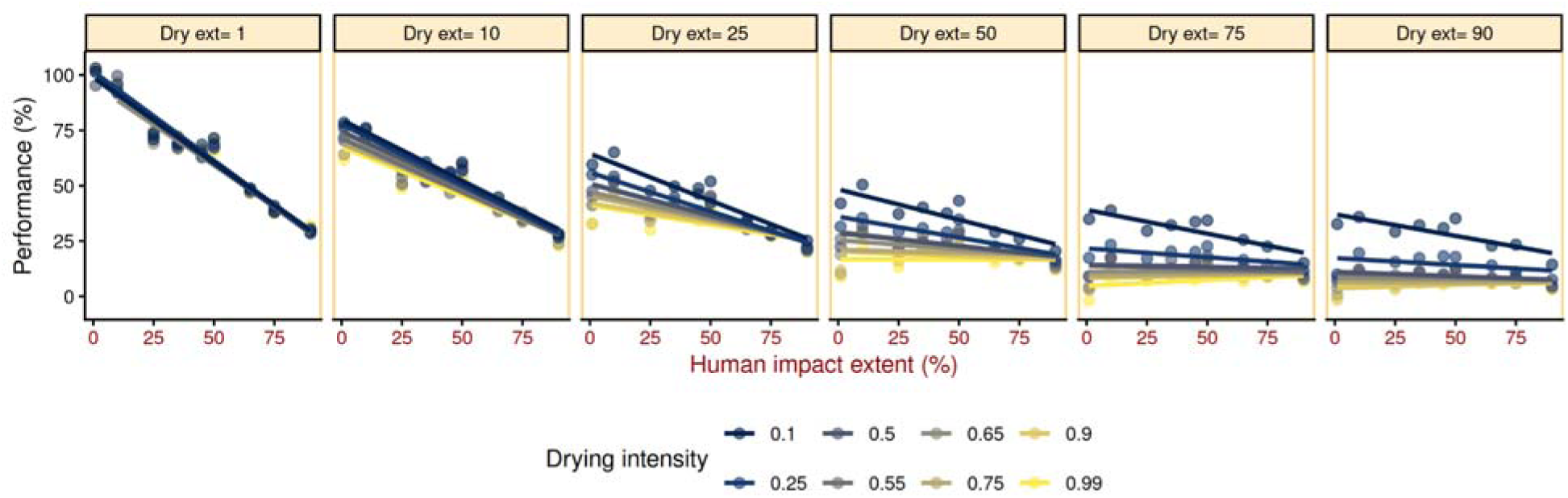
Linear relationship between biological index performance (%) and human impact extent (% of impacted sites) along selected drying extent scenarios and for different drying intensities (blue to yellow lines).

### Coupling real drying patterns with simulated scenarios

Drying extent and intensity from six non-impacted drying river networks aligned well with the drying scenarios that we designed for the synthetic river network (Figure 5). In these river networks, drying extent (i.e., number of sites that dry) ranged from low to almost fully drying networks (Lepsämänjoki, “A” = 30.30%, Bükkösdi, “B” = 42.30%, Albarine “C” = 55.53%, Butižnica, “D” = 55.63%, Velička, “E” = 64.23%, and Genal, “F” = 84.41%). Similarly, drying intensity (i.e., the variability in drying length for all network sites) ranged from low to very high (A= 0.25, B= 0.65, C= 0.50, D= 0.99, E= 0.50, and F= 0.9). Under conditions of 10% human impact extent (Figure 5, see Figure S8 for the other %), the performance of the biological index rapidly decreased as a function of drying extent and intensity. Each river network showed different performance profiles, with the Genal river network (F) being the most extreme case, where performance was below 10% (Figure 5). Thus, for this river network, the capacity of the biological index to distinguish between impacted and non-impacted sites would be reduced by 90% solely due to its natural drying conditions.

**Figure 5.**
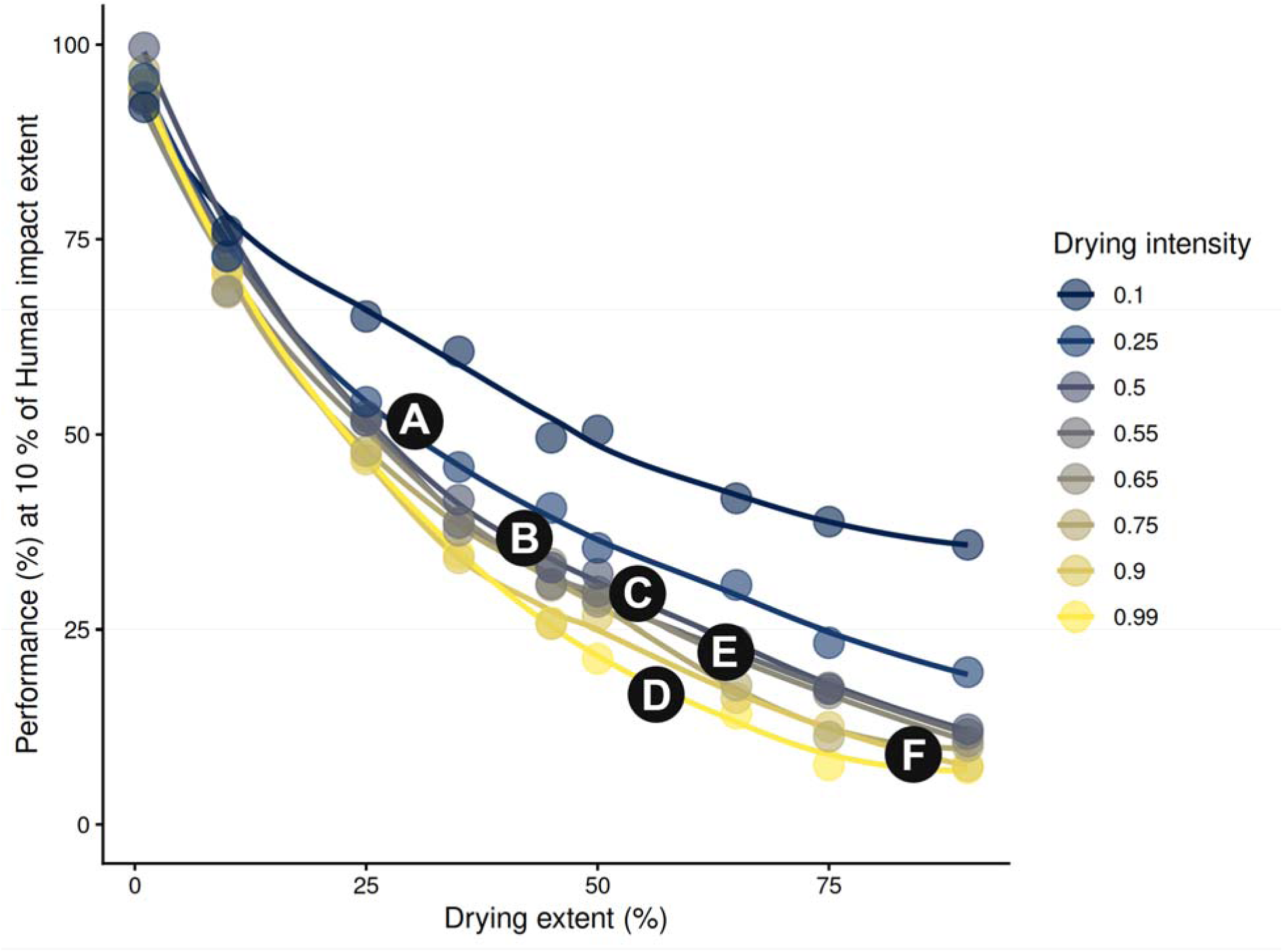
Biological index performance along the drying extent gradient (%) and for different drying intensities (coloured lines and dots) under a 10 % human impact extent. Dark circles indicate the position that would correspond to the six low-impact European drying river networks according to their drying extent and intensity (A-F). From left to right, Lepsämäanjoki catchment, Finland (“A”), Bükkösdi catchment, Hungary (“B”), Albarine catchment, France (“C)”, Butižnica catchment, Croatia (“D”), Velička catchment, Czechia (E), and Genal catchment, Spain (F).

## Discussion

Our simulations revealed the multifaceted effects of drying on diversity and biological index performance across spatial and temporal scales (i.e., extent and intensity). In line with our first hypothesis, increased fragmentation of the river network resulted in lower local richness and scores of the biological index, which is consistent with previous studies (Chalmandrier et al., 2025; Crabot et al., 2020; Escobar Camacho et al., 2025; Journiac et al., 2025). While drying extent acted as a regional modulator defining the maximum local richness (Cunillera Montcusí et al., 2025; Riva & Fahrig, 2023; Zhang et al., 2024), drying intensity defined the rate at which local richness was lost in response to drying-driven fragmentation, which increased as the intensity increased. This trend aligns with previous regional and large-scale studies that showed how drying duration and frequency diminish local diversity (Arias Real et al., 2021; Escobar Camacho et al., 2025; Pineda-Morante et al., 2022). In line with our second and third hypotheses, index performance drastically dropped as soon as drying started, with a 50% decrease under a 25% drying extent. Then this trend leveled off as drying extent continued to increase, with a decrease of around 75% in performance above 50% of the drying extent. Above this extent, impacted and non-impacted sites would be similarly classified by the biological index, leading to the failure of any monitoring assessment. Thus, our results show that, at a certain drying extent, biological indices would fail to capture changes in the biological quality associated with human impacts. Such insights are essential to better frame monitoring practices in drying river networks (Cid et al., 2020; Datry et al., 2023).

Focusing only on local niche-based approaches constitutes a weakness for traditional biological indices when regional-scale processes such as species dispersal can be affected by changes in river network connectivity resulting from drying (Courtwright & Hawkins, 2024; Haase et al., 2023). Therefore, a metacommunity perspective has been proposed to integrate local and regional scales (Cid et al., 2020; Chase et al., 2020). Yet, spatiotemporal connectivity of drying river networks can be diverse and generate a wide variety of configurations (Datry et al., 2023; Journiac et al., 2025; Sarremejane, Silverthorn, et al., 2024). This inherent complexity, in turn impacts biological index performance, which changes greatly depending on drying extent or intensity. For example, while 50% drying extent implied a 50% drop in performance in low drying intensity scenarios, the same drying extent in high drying intensity scenarios resulted in a 75% drop in performance. Our approach can cover a wide range of drying scenarios and adapt to real drying river networks when surface water permanence information is available (Messager et al., 2021; Mimeau et al., 2025). Therefore, rather than providing a unique answer to the question of “when will biological indices fail?”, our approach can measure the expected degree of failure of a specific biological index along different gradients of stress. This logic builds on the method proposed by Cid et al. (2020), which used biological traits, regional connectivity, and flow regime in a model to predict index performance. Here, we advanced these ideas by using simulations of spatiotemporal connectivity to capture regional-scale variability of these highly dynamic ecosystems (Cunillera Montcusí et al., 2023; Datry et al., 2016; Journiac et al., 2025).

Our model confirms the predictions suggested by Cid et al. (2020), showing that the index performance follows a convex decay in performance, which became more negative as drying intensity increased. This decay pattern in index performance is caused by the fragmentation of the network, which reduces dispersal between habitats (Fournier et al., 2023; Sarremejane, Silverthorn, et al., 2024), but also by the increased homogenisation of communities generated by the dominance of species able to survive in these scenarios (Cid et al., 2020; He et al., 2020; Heino et al., 2017). These two drivers were captured by our simulations through 1) the concomitant decrease in spatiotemporal connectivity, measured as the increase in dispersal resistance between habitats, and 2) the homogenisation linked to human impact extent that favoured the colonisation of tolerant taxa (Alba-Tercedor et al., 2002). This was reflected in the progressive drop in performance as human impact extent increased, concomitantly increasing the similarity between impacted and non-impacted communities due to mass effects (i.e., increased presence of tolerant taxa across the network led to their dominance also in non-impacted habitats through dispersal). Although the decrease in the performance of the biological index when human impacts were widespread may seem counterintuitive, from a metacommunity standpoint, widespread human impacts may favour homogenisation even in non-impacted sites. This is because of greater dispersal pressure from impacted sites, which resulted from source-sink dynamics (Horváth et al., 2025; Leibold & Chase, 2017; Savary et al., 2024). Indeed, these results were related to the species pool of our model, which was mostly composed of weaker dispersers that use the stream network as their main dispersal route (62% of the species pool), which corresponded to a convex decay pattern (Cid et al., 2020). Therefore, here we showed that the use of biological indices not adapted to drying underestimates true biological quality as soon as a small percentage of the catchment dries (i.e., drop in 25% in performance with only 10% of the reaches drying). Such insights can benefit decision making by determining the validity of biological indices and decide what alternative approach could prove more effective (Courtwright & Hawkins, 2024; Steward et al., 2022; Wilding et al., 2018).

Drying river networks are still poorly managed in most cases, so novel biomonitoring approaches that capture spatiotemporal dynamics are urgently needed (Datry et al., 2016, 2023; Stubbington et al., 2024). This could be done by using non-impacted drying river networks as reference conditions and correcting for the drying effect on ecological quality assessment. However, categorizing drying river networks into defined compartments is risky, as it may only partially capture the complexity of these ecosystems (Chadd et al., 2017; Wilding et al., 2018). Such an oversimplification could, in turn, lead to a similar decrease in performance when conditions diverge from the defined reference (as in Figure S7). Other options include the use of new indices that incorporate terrestrial fauna inhabiting the dry riverbed (Bruno et al., 2022; Sánchez-Montoya et al., 2020; Sánchez-Nogueras et al., 2025; Stubbington et al., 2019) or fauna inhabiting river pools during the dry season (Ersoy et al., 2024). When catchment-level drying patterns are available (e.g., Mimeau et al., 2024), our approach could be used to simulate case-specific expected drop in performance (Blair, 2021; De Koning et al., 2023). This would provide a more individualised river network assessment of index performance that could help identify sites or scenarios with lower performance (Blair, 2021; Juarez et al., 2021). Therefore, the support provided by simulations could help to integrate the natural complexity of drying river networks and facilitate a more realistic assessment leading to improved monitoring, restoration and conservation of these ecosystems (Datry et al., 2023; Stubbington et al., 2022).

In this study, we used a metacommunity model that simulates the exchange of individuals between habitats following a determined network structure (Borthagaray, Cunillera-Montcusí, Bou, Tornero, et al., 2023; Cunillera Montcusí et al., 2021). While such an approach has proven useful for capturing spatially driven assembly patterns (Borthagaray, Cunillera-Montcusí, Bou, Biggs, et al., 2023; Cunillera Montcusí et al., 2025), it simplifies reality by neglecting biotic interactions (Alahuhta et al., 2025; McIntosh et al., 2017), topographic and land use-related landscape barriers (Cañedo Argüelles et al., 2015; Firmiano et al., 2021), or drying resistance traits (Cid et al., 2020; Sarremejane et al., 2021; Tolonen et al., 2019). In this regard, Cid et al., (2020) hypothesized that resistance to drying would lead to different decay trends in performance. Our simulation did not include such traits, which could be included assuming that some species remain unaffected when a site dries. However, the lack of robust data on resistance traits and the interaction of these traits with human impacts (i.e., biological index scores) was beyond the scope of this work and deserves further study. For example, the role of resistance strategies (Doretto et al., 2020; Hershkovitz & Gasith, 2013; Strachan et al., 2014) and the influence of community structure prior to drying, also known as community disassembly (O’Neill, 2016), are interesting topics for future studies. In contrast, the availability of biological indices that quantify the degree of tolerance to human impacts allowed us to explore the interaction between drying and human impacts. This information enhanced the realism of the simulations by incorporating key functional aspects in response to local-scale stressors, which in turn interacted with different dispersal strategies reflecting organisms’ regional-scale responses (Cunillera Montcusí et al., 2021; Jacquet et al., 2022).

Despite the limitations mentioned, the current approach improved our understanding of the interactive effects of natural drying and human impacts on the performance of the biological index for both simulated and real drying river networks. While previous simulations explored the interplay between drying and diversity in drying river networks (Jacquet et al., 2022; Journiac et al., 2025), our work adds a novel layer of information by incorporating human impacts (Alba-Tercedor et al., 2002). In this line, our approach could easily be adapted to other aquatic organisms such as diatoms, fish, or terrestrial invertebrates from dry riverbed (Sánchez Montoya et al., 2016; Sánchez-Montoya et al., 2020) by considering their dispersal capacities, ecological niche and any other ecological characteristic that determines their responses to human impacts. The combination of these simulations would advance towards more complex digital tools, able to realistically simulate ecosystem dynamics (De Koning et al., 2023).

Overall, we assessed the role that drying plays in altering biological index performance by modelling metacommunity dynamics –i.e., dispersal limitation– as well as family-based functional traits –i.e., dispersal strategies and tolerance. In this sense, our approach represents a foundational step towards adjusting and/or developing novel models specifically acknowledging dynamics of drying river network to challenge niche-based approaches (Chase et al., 2020; Cid et al., 2020; Datry et al., 2023). The increasing availability of high-resolution data from new technologies and citizen science can provide a more accurate vision of spatiotemporal connectivity in these dynamic systems, allowing catchment-specific assessments (Fernández Calero et al., 2024; Pineda-Morante et al., 2022; Truchy et al., 2023). Such high spatiotemporal resolution enables the development of more realistic simulations for drying river network, as demonstrated in this study. These models can support decision-making and future scenario simulations while accounting for fluvial metacommunity dynamics (Cid et al., 2020; Chase et al., 2020). The full potential of these tools remains to be unlocked, but they may prove essential for addressing future ecological and conservation challenges (De Koning et al., 2023; Maasri et al., 2022), helping ecologists and managers break current conservation halts (Haase et al., 2023).

## Acknowledgments

This study is funded by the project “DRY-Guadalmed: Herramientas avanzadas para la evaluación del estado ecológico de ríos temporales mediterráneos durante la fase seca” (PID2021-126143OB-C21 and PID2021-126143OB-C22), and partly by the DRYvER project (Horizon 2020 #869226). This research was carried out in the FEHM (Freshwater Ecology, Hydrology and Management) research group funded by the “Agència de Gestió d’Ajuts Universitaris i de Recerca” (AGAUR) at the “Generalitat de Catalunya” (2021SGR00692). Additional funding was also received from the “ERDF A way of making Europe”. NB is a Serra Húnter fellow and is supported by an ICREA Academia 2021 award from the Catalan Institution for Research and Advanced Studies. ZE is also supported by the Programa Atracción del Talento (Comunidad de Madrid, No: 2024-T1/ECO-31557). DCM is also supported by the Research Executive Agency research and innovation program under Marie Sklodowska-Curie Grant Agreement No. 101062388. Since 16 April 2025, MCA has been seconded to the ERC Executive agency. The views expressed in this paper are purely those of the author. They do not necessarily reflect the views or official positions of the European Commission, the ERC Executive Agency or the ERC Scientific Council.

## Author contributions

Zeynep Ersoy, David Cunillera-Montcusí, Núria Bonada and Núria Cid conceived the ideas and designed the conceptual framework; Zeynep Ersoy and David Cunillera-Montcusí collected the data for simulations from the literature (equal contribution), analyzed the data (equal contribution) and led the writing of the manuscript (equal contribution). Núria Bonada and Núria Cid provided supervision and critical review. Núria Bonada, Núria Cid and María Mar Sánchez-Montoya led the project funding this work. All authors contributed critically to the drafts and gave final approval for publication.

