## Appendix S1 for "Simulating drying and human impacts on river networks to evaluate biological quality indices performance through the lens of metacommunity theory"

**Supplementary Information**

**Appendix S1:** In the following lines, we define in detail the metacommunity model that we used for each simulation and its parameterization. This model has already been used in different publications, from which other detailed information can be obtained (Cunillera-Montcusí et al., 2021, 2025; Borthagaray et al., 2025)

**Methods for the metacommunity simulations**

The metacommunity model used in this study aimed to capture diversity patterns considering major regional heterogeneity drivers (i.e., Drying extent, Drying intensity, and Human impacts). For each combination of these factors, we ran 10 simulations. Each simulation generated a metacommunity (sites X species matrix) from which we later calculated taxon richness –representing river invertebrate families– and biological index values according to family tolerance scores.

*1. 1. Metacommunity model parameters*

All coalescent model runs had the same parameters (i.e., species pool, number of species with a determined dispersal ability) except for drying and human impacts. Drying affected community size (i.e., surrogate of community total abundance) and dispersal connectivity (i.e., STconmat values), while human impact extent determined local establishment penalty based on family tolerances. To ensure a realistic species pool, we compiled all taxa (n= 120) used within the biological index (Alba-Tercedor et al., 2002) and their human impact tolerance values (see Table S1). Different dispersal abilities (drifters, swimmers/crawlers, and flyers) were assigned for each family based on their dispersal strategy traits (aquatic passive, aquatic active, aerial active), which are described using a fuzzy-coding approach from the DISPERSE database (Sarremejane et al., 2020). Because the DISPERSE database used the genus level, we averaged the trait values of all genera within each taxon used in the biological index. If the value for each dispersal category was higher than the mean, we used that dispersal category; otherwise, the same taxon was repeated with different dispersal strategies. For example, if a taxon was both a flyer and a drifter, we considered it as 2 taxa (1 flyer and 1 drifter), which makes up a total of 182 taxa in the regional species pool (see Table S1).

Drying defined two key metacommunity parameters:

First, community size (J), which represented the number of individuals a node could sustain. Community size was determined as a function of drying frequency, where fully perennial nodes contained a maximum of 200 individuals (J_max_). Then, the number of individuals in each site was defined as J_i_=(-J_min_+(J_max_/(Max_Permanence^0.8^))*Permanence_i_^0.8^), where J_max_= 200, Permanence_j_ is the number of flow days (days with no drying) for site *i* and Max_Permanence is the maximum value of permanence that each node could have (i.e., 48). Note that the term individual in the model refers to an aggregation of individuals from the same taxa (e.g., local populations), not a single organism.

Second, drying defined migration probability based on three connectivity matrices, each representing the three dispersal groups (i.e., STconmat for drifters, swimmers, and flyers). These three matrices quantified migration probabilities as a function of both distance and drying conditions for each dispersal group and all three were incorporated together within the same simulation (i.e., 61 drifter, 51 swimmer/crawler, and 70 flyer taxa; Cunillera-Montcusí et al., 2021). We incorporated resource overlap between dispersal groups, considering half of the available community size (J) to be shared among all dispersal groups, and the remaining half to be exclusive to each group ( ⅓ per group). We did this to avoid dispersal-driven exclusion of other dispersal groups (i.e., drifters and swimmers) by aerial active dispersers, which are not restricted to dendritic connectivity. With this, we ensured the presence of drifters and swimmers in more isolated communities (Cunillera-Montcusí et al., 2021). Therefore, within each community, the shared part of its size (J/2) could be occupied by taxa belonging to any dispersal group, while each third of the remaining part could only be occupied by taxa from a specific dispersal group.

Human impact was modeled as a local penalty in sites that were defined as “Impacted”. This penalty reduced the establishment probability of taxa based on their classification within the biological index (Alba-Tercedor et al., 2002). This index assigns each taxon a score between 1 (highly tolerant to impact) to 10 (highly sensitive to impact). To apply this in the metacommunity model, we used the inverse of these scores to define an environmental impact penalty, which modified the establishment probability of each taxon in each site. This penalty ranged from 0.01 for highly sensitive taxa (reducing their establishment probability to 1% of its value defined by migration probability) to 0.91 for highly tolerant taxa (reducing the establishment probability to 91% of its value defined by migration probability). These penalties were assigned independently of the drying pattern. In sites experiencing both drying and human impact, the penalty strength was adjusted, decreasing it for sensitive taxa and increasing it for tolerant taxa as a function of drying. In contrast, sites assigned as “Non-impacted” had no penalty assigned, implying that taxa establishment probability was only determined by migration probabilities.

*1. 2. Metacommunity model coalescent runs*

We simulated the coalescent filling of each metacommunity 10 times for each drying and human impact combination. This simulation consisted of an iterative filling of all communities (i.e., sites) based on the probabilities defined by a migration matrix (M), which was in turn modified by human impact-based penalties. We assumed that there was no arrival of species from other river networks. Each simulation began with randomly adding one individual from the species pool to each community. Subsequently, each local community was filled to its size (J) by sequentially sampling individuals from three sources: 1) species pool (*m.pool*), 2) neighboring communities dispersal (*m.neighbor*), or 3) self-recruitment. We set the *m.pool* parameter to 0.01 in all scenarios to define a system where dispersal and self-recruitment were the main sources of diversity. Migration from neighboring communities (*m.neighbor*) was defined by a kernel based on an exponential decay function of STconmat values. STconmat defined dispersal resistance combining distance and drying-induced fragmentation (Cunillera-Montcusí et al. 2023). The migration probability (*m_ij_*) between two communities i and j was calculated as: *m_ij_ = m.max ∗ e^−b∗STconmatij^*, where *STconmat*_ij_ is the STconmat value between communities i and j, and *m.max* was set to 1, representing the maximum migration rate between communities at zero distance (Botharagay et al., 2023; Borthagaray et al., 2025). The parameter *b* determines the rate at which migration probability decays with STconmat, here defined as the value at which this probability decays to half of its maximum, as b = −log(0.5) / d_50_. We set d_50_ as 0.15 for all simulations to define a constant rate of decay for all dispersal groups, which were already submitted to different STconmat dispersal resistance probabilities (drifters, swimmers, and flyers). Finally, we calculated self-recruitment probability based on the difference, considering the two previous values as 1− (m.pool + m.neighbor). Based on these probabilities, we defined a community-by-community migration matrix (M). These matrix probabilities were then modified according to the penalties defined within the corresponding human impact scenario. Once all communities were filled (J), we quantified local diversity (i.e., alpha diversity) and the biological index by summing all scores of families found in each community (i.e., site-level biological quality). For each scenario combining drying and human impact, we calculated the median across the 10 replicates of these metrics to obtain an averaged value for the focal scenario.

**Supplementary plots:**

**A)**

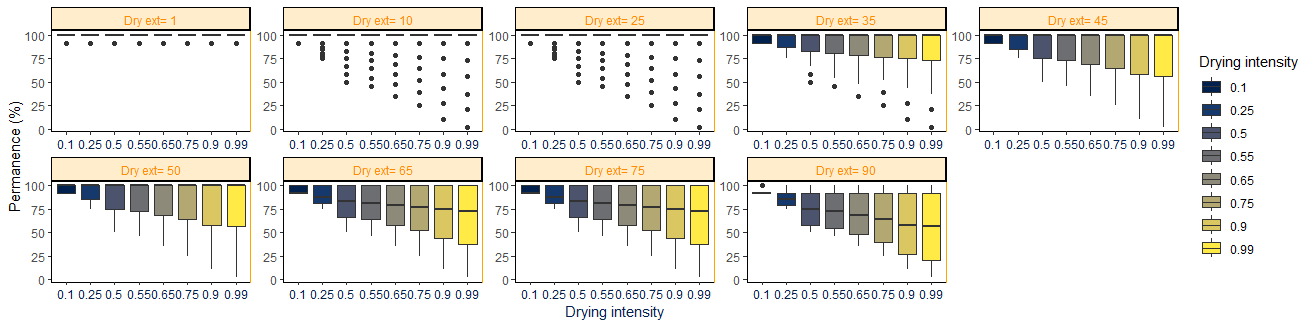

**B)**

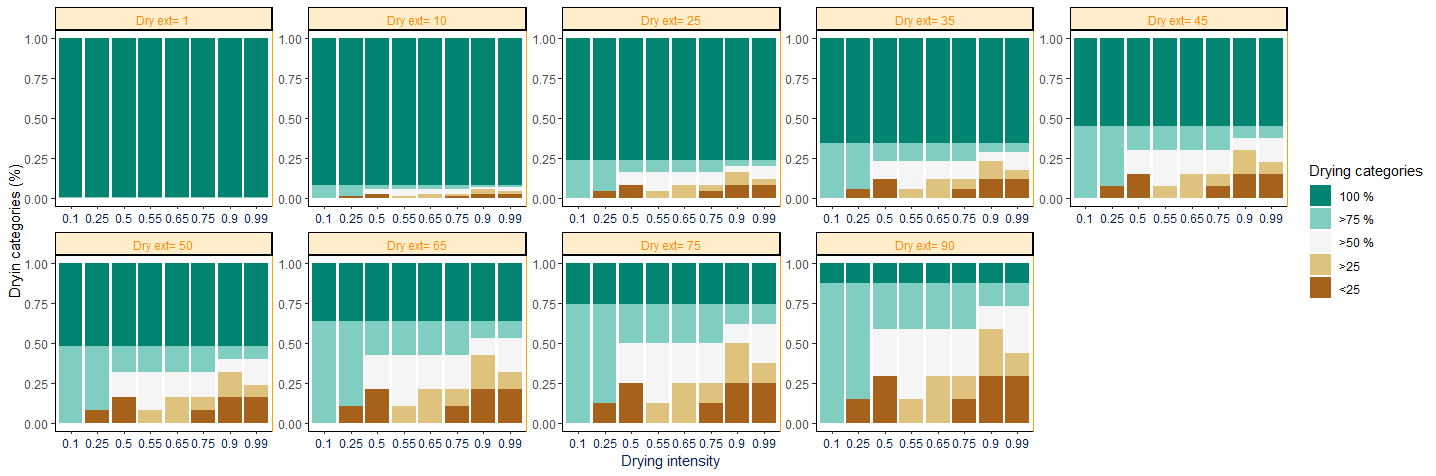

**C)**
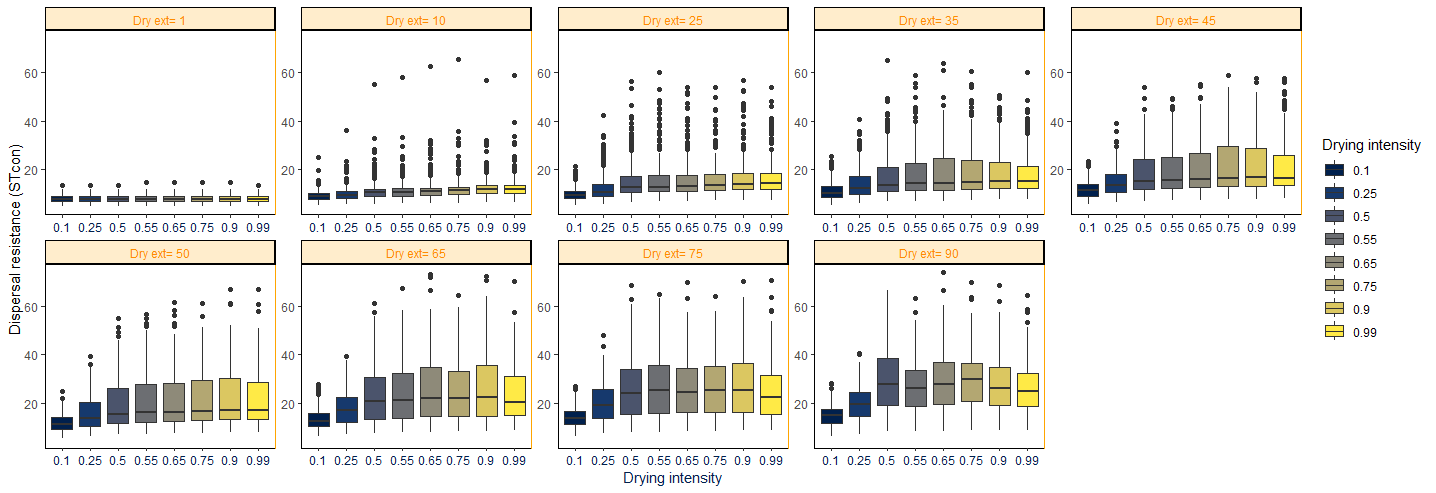

**Figure S1.** A) Summary results for each combination of drying extent (each panel) and intensity scenarios (x-axis). A) Distribution of sites’ permanence. B) Distribution of sites’ drying categories. C) Distribution of sites’ STconmat (i.e., dispersal resistance based on distance and drying isolation). Drying intensity is shown in each boxplot colour (blue to yellow colours ranging from 0.1 to 0.99). Drying categories are shown in barplot colour (green to brown colors ranging from 100 % to <25 %).

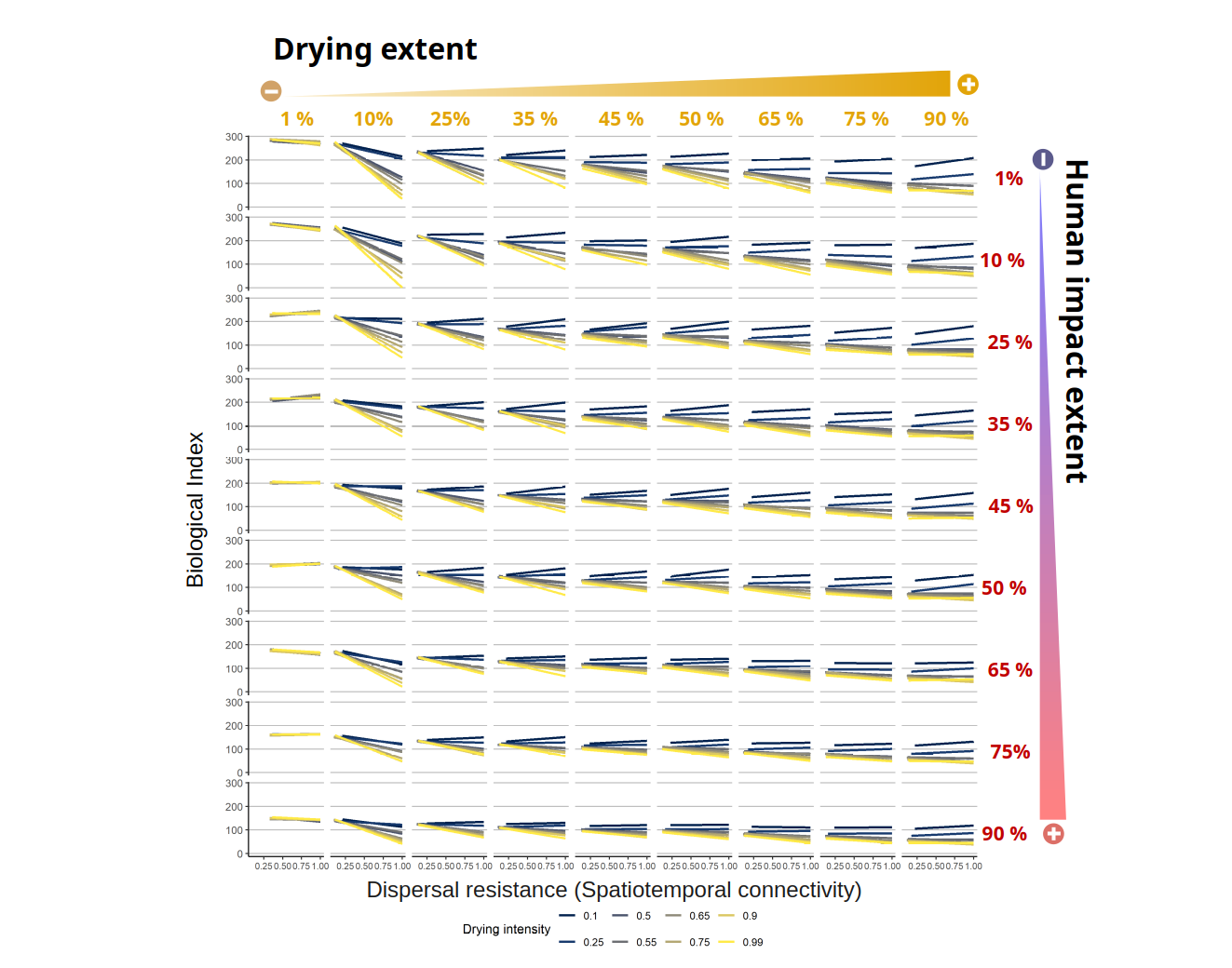

**Figure S2.** Linear relationships between biological index (y axis) and dispersal resistance (i.e., spatiotemporal connectivity calculated as STconmat; x axis). Each panel shows a different combination of scenarios along the human impact extent (vertical axis) and drying extent (horizontal axis). Drying intensity is shown in each smoothing line colour (blue to yellow colours ranging from 0.1 to 0.99). Points have been removed to facilitate interpretation.

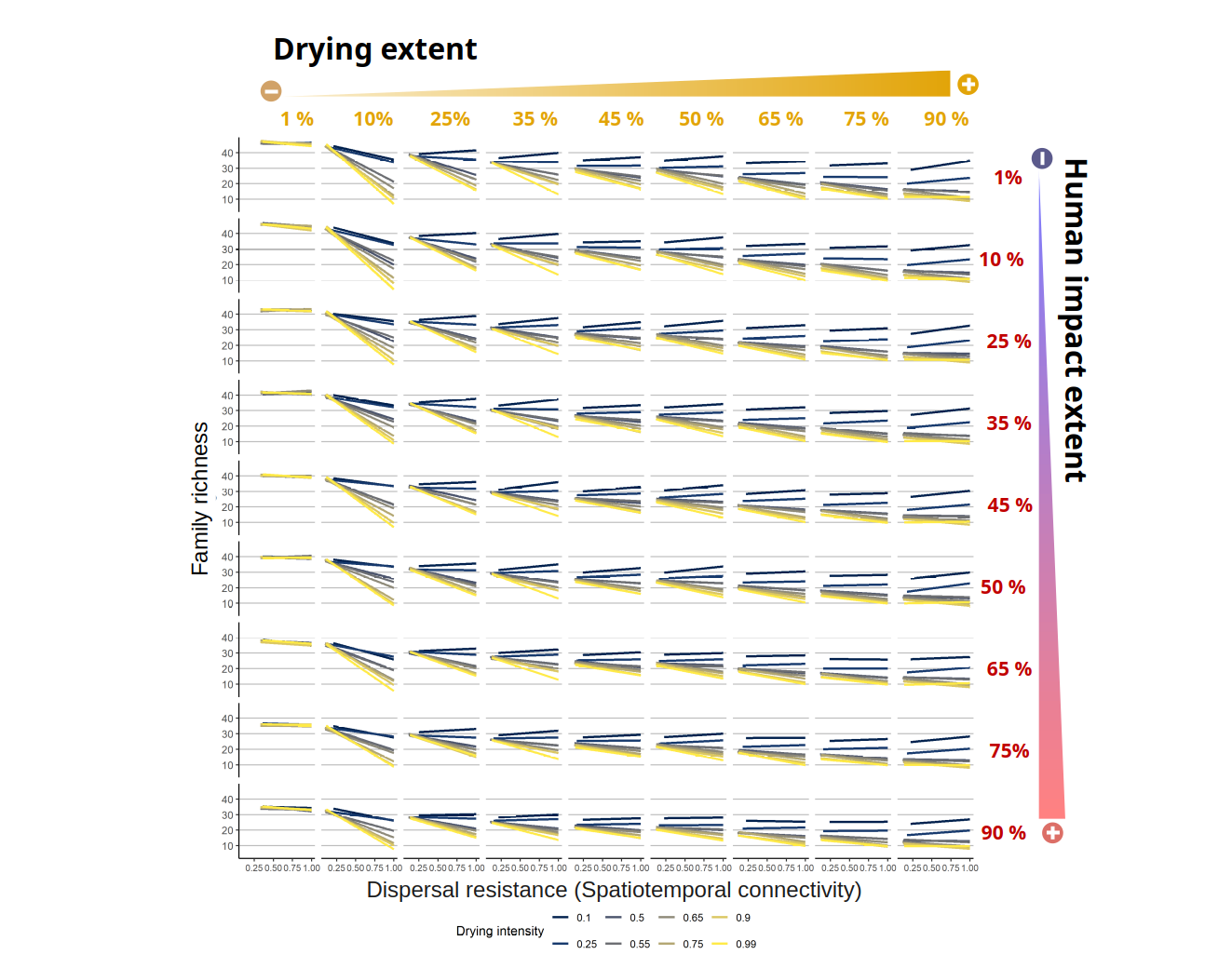

**Figure S3.** Total relationships between family richness (y axis) and dispersal resistance (i.e., spatiotemporal connectivity calculated as STconmat; x axis). Each panel shows a different combination of scenarios along the human impact extent (vertical axis) and drying extent (horizontal axis). Drying intensity is shown in each smoothing line colour (blue to yellow colours ranging from 0.1 to 0.99). Points have been removed to facilitate interpretation.

**Plot A)**
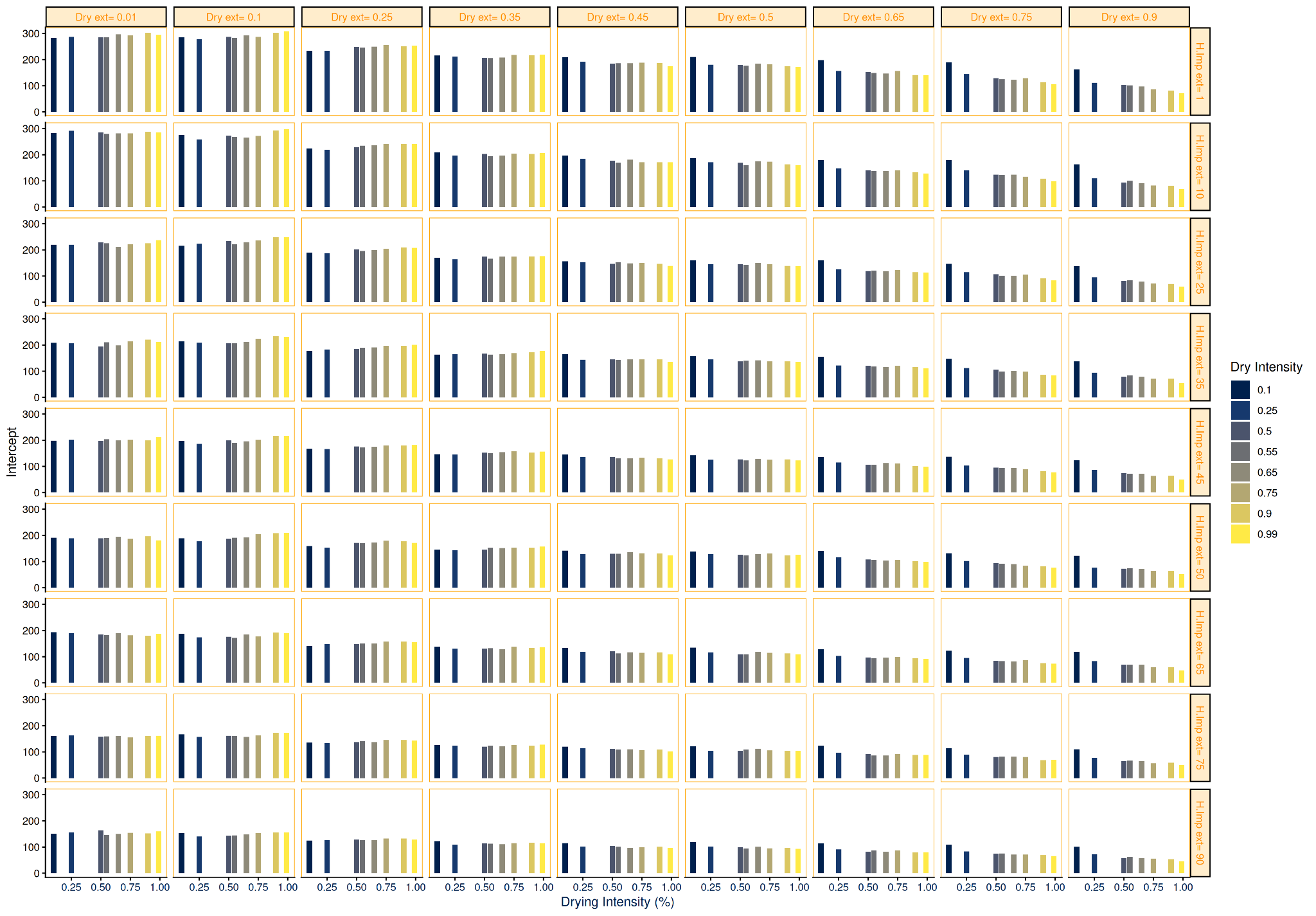

**Plot B)**

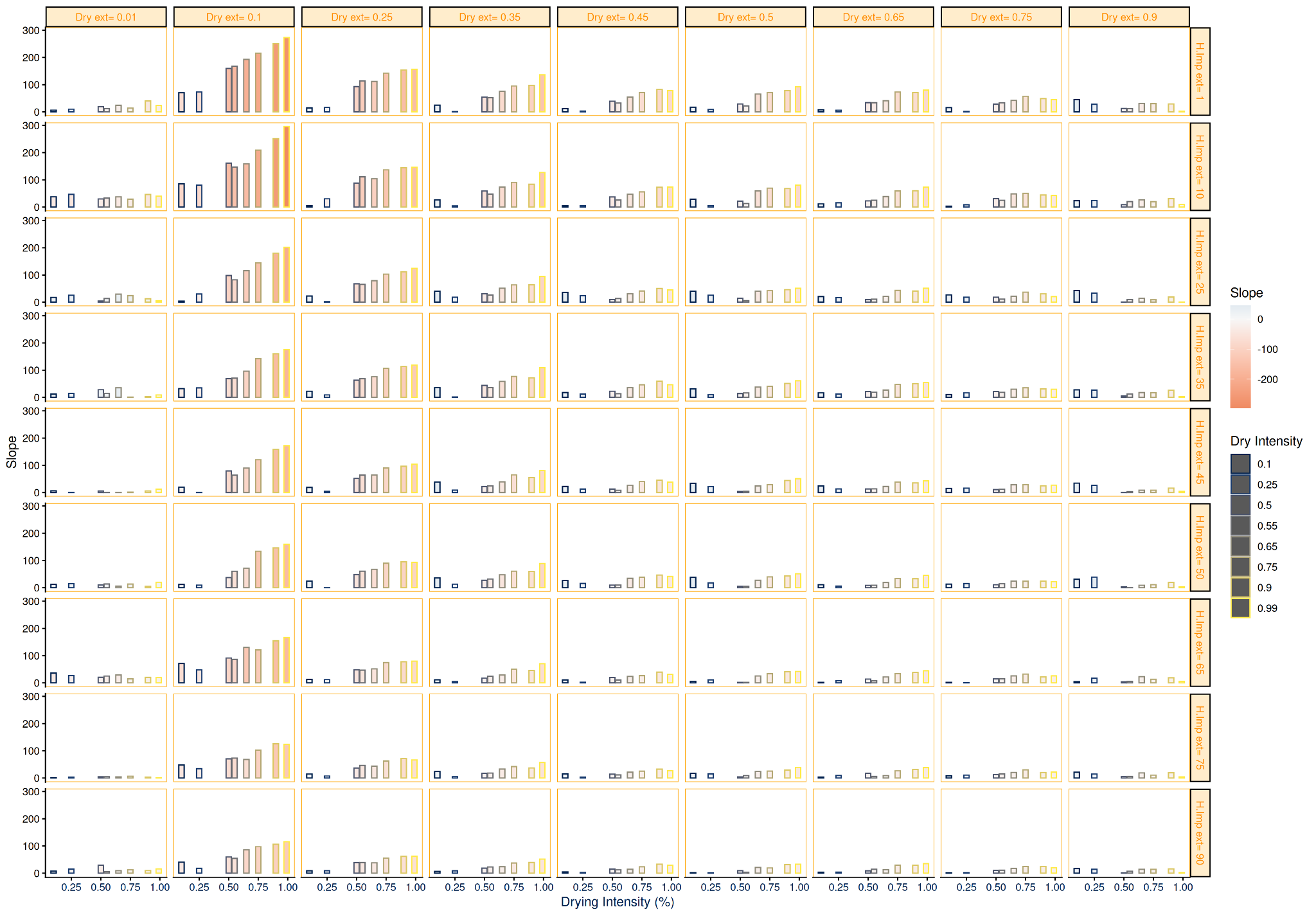

**Figure S4.** Summary results of the intercept and slope from the relationship between biological index and STconmat (i.e., dispersal resistance based on distance and drying isolation) for all combinations of scenarios. Each panel shows the different combinations of scenarios with human impact extent (y axis), drying intensity (x axis) and drying extent corresponding to each facet column (ranging from 0.1 to 0.99). Plot A indicates the model intercept values with point size (Biological index value when STconmat = 0). Plot B represents model slopes with point size (absolute slope value) and point filling colour (blue colours being positive and red colours being negative).

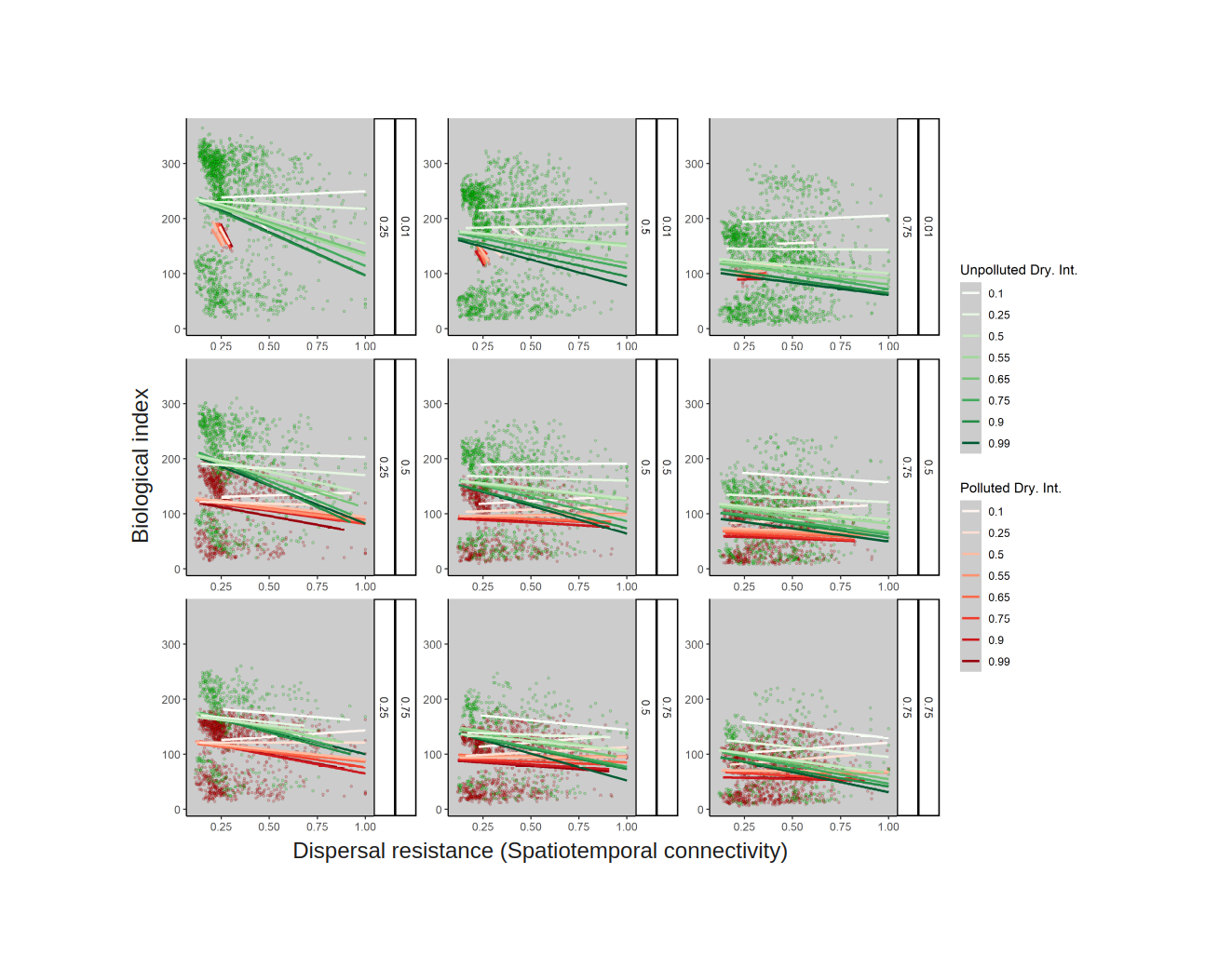

**Figure S5.** Linear relationship between biological index and dispersal resistance (i.e., spatiotemporal connectivity calculated as STconmat) across selected scenarios of drying extent (25 %, 40 %, 75 %), and human impact extent (1 %, 50 %, 75 %) at the whole gradient of drying intensity. Coloured points (green and red) indicate human impact categories (unimpacted and impacted, respectively). Coloured lines indicate the different levels of drying intensity for each human impact category.

**Figure S6.** Linear relationship between biological index performance (%) and human impact extent (%) across all drying extent scenarios. Drying intensity is shown in each smoothing line colour (blue to yellow colours ranging from 0.1 to 0.99). The reference state is unique for all the scenarios and corresponds to a pristine river (Human impact extent = 0.01) with low drying extent (Dry ext = 0.01) and low drying intensity (Dry intensity = 0.1).
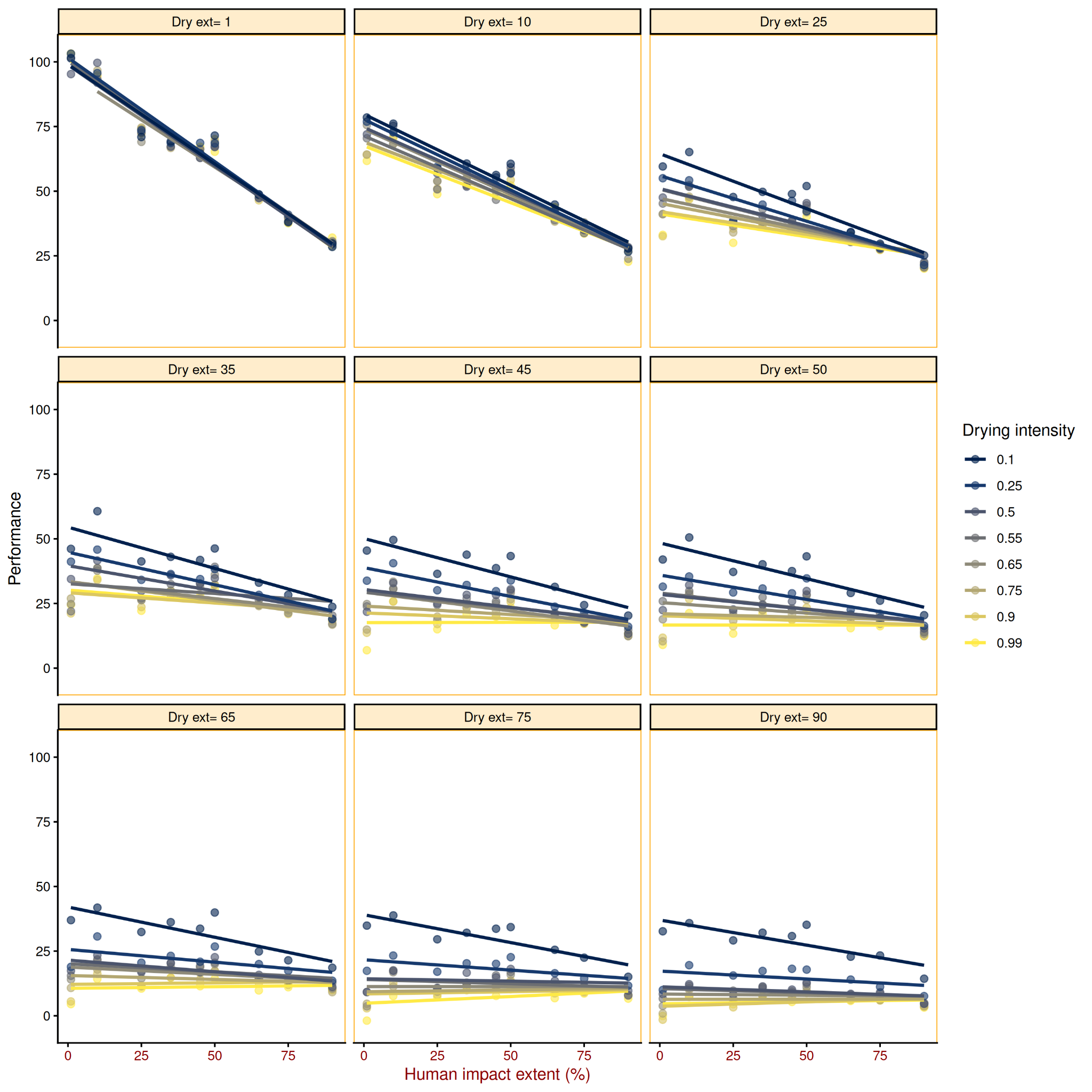

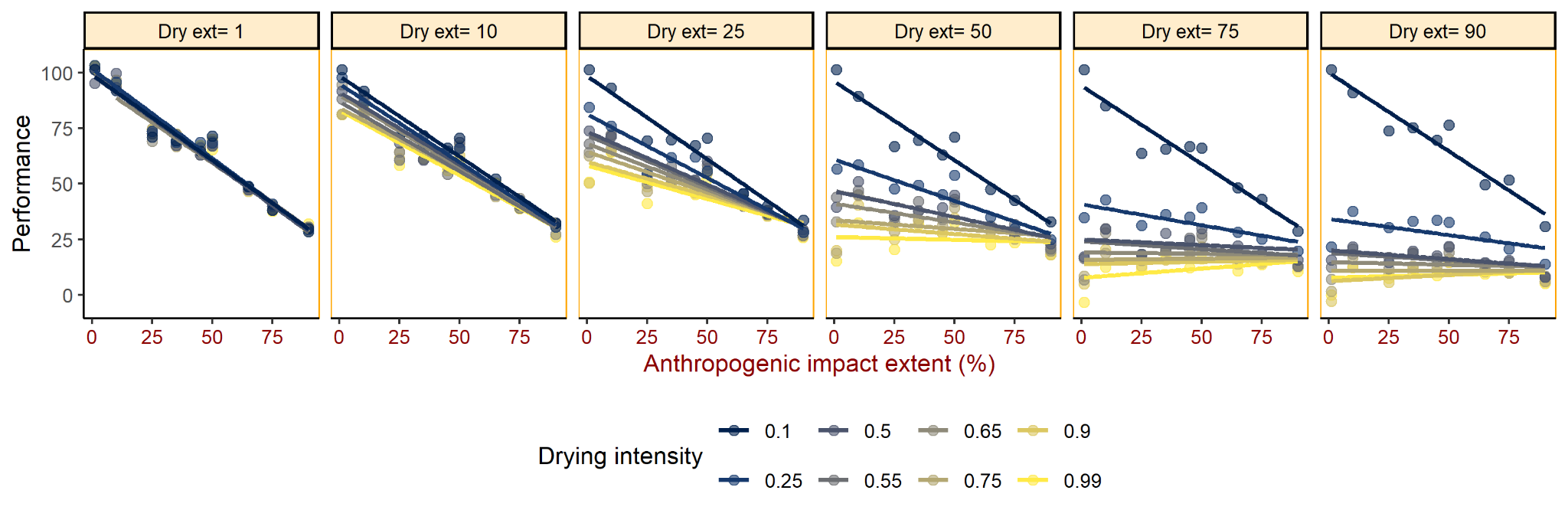

**Figure S7.** Linear relationship between the biological index performance (%) and human impact extent (%) along selected drying extent scenarios (1, 10, 25, 50, 75 and 90%). Drying intensity is shown in each smoothing line colour (blue to yellow colours ranging from 0.1 to 0.99). The reference state is set for each drying extent scenario and corresponds to an unimpacted river (Human impact extent = 0.01) with low drying intensity (Drying intensity = 0.1).

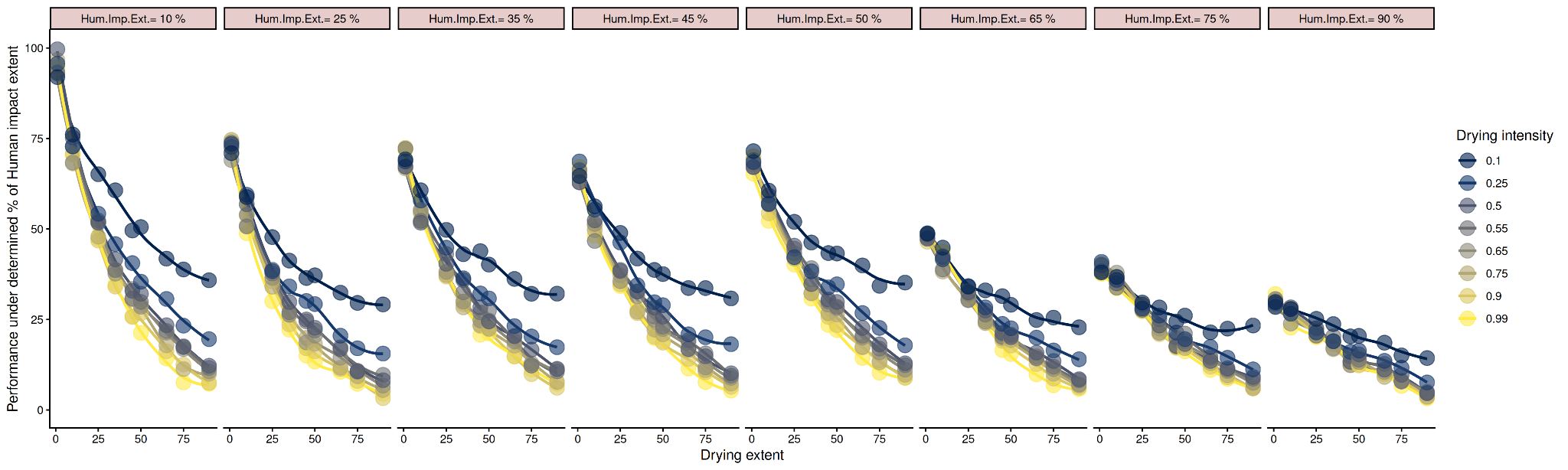

**Figure S8**: Biological index performance (%) along the drying extent gradient (%) and for different drying intensities (colored lines and dots) along the whole gradient of human impact extent (horizontal panels).
